# Elevated intracellular cAMP concentration mediates growth suppression in glioma cells

**DOI:** 10.1101/718601

**Authors:** Dewi Safitri, Harriet Potter, Matthew Harris, Ian Winfield, Liliya Kopanitsa, Ho Yan Yeung, Fredrik Svensson, Taufiq Rahman, Matthew T Harper, David Bailey, Graham Ladds

**Author notes:** Corresponding author: Dr Graham Ladds, Department of Pharmacology, University of Cambridge, Tennis Court Road, Cambridge, CB2 1PD.

## Abstract

Supressed levels of intracellular cAMP have been associated with malignancy. Thus, elevating cAMP through activation of adenylyl cyclase (AC) or by inhibition of phosphodiesterase (PDE) may be therapeutically beneficial. Here, we demonstrate that elevated cAMP levels suppress growth in C6 cells (a model of glioma) through treatment with forskolin, an AC activator, or a range of small molecule PDE inhibitors with differing selectivity profiles. Forskolin suppressed cell growth in a protein kinase A (PKA)-dependent manner by inducing a G_2_/M phase cell cycle arrest. In contrast, trequinsin (a non-selective PDE2/3/7 inhibitor), not only inhibited cell growth via PKA, but also stimulated (independent of PKA) caspase-3/-7 and induced an aneuploidy phenotype. Interestingly, a cocktail of individual PDE 2,3,7 inhibitors suppressed cell growth in a manner analogous to forskolin but not trequinsin. Finally, we demonstrate that concomitant targeting of both AC and PDEs synergistically elevated intracellular cAMP levels thereby potentiating their antiproliferative actions.

## 1. INTRODUCTION

Glioma is a general term for brain tumours that originate from glial cells in the central nervous system and which may progressively lead to death if not treated early (Mamelak and Jacoby 2007, Harris et al. 2018), with glioblastoma, the most common type of glioma, showing particularly poor survival (Schwartzbaum et al. 2006). The development of novel therapeutic approaches targeting glioma and glioblastoma are urgently required.

Defects in a number of signalling pathways have been reported in glioma pathogenesis, including the phosphatidylinositol-3 kinases/ phosphatase and tensin/ protein kinase B/ mammalian target of rapamycin (PI3K/PTEN/Akt/mTOR) cascade; the retinoblastoma pathway (pRB); the Ras/ mitogen-activated protein kinase (RAS/MAPK) pathway; signal transducer and activator of transcription 3 (STAT3); zinc transporter 4 (ZIP4); as well as the adenylyl cyclase (AC) system (Li et al. 2016, Burns et al. 1998, Holland et al. 2000, Mao et al. 2012).

Importantly, lower cAMP levels are observed in brain tumour tissue (25.8 pmoles/mg) compared to normal healthy tissue (98.8 pmoles/mg) (Furman and Shulman 1977). Indeed forskolin, an AC activator that elevates cAMP levels, has shown promising results in ameliorating cancer development (Sapio et al. 2017, Illiano et al. 2018). While high levels of intracellular cAMP may kill cancer cells, exact molecular mechanisms have not been clearly elucidated. Direct elevation of cAMP using the cAMP analogues 8-bromo-cAMP, 8-chloro-cAMP, monobutyryl cAMP and dibutyryl cAMP, however, cannot be recommended due to the toxicity of these compounds (Hirsh et al. 2004). Thus, there is a need to develop safe and effective compounds that can increase cAMP levels by pharmacological intervention.

Depending on the cell type where the malfunctions are observed cAMP may play a role in either promoting or suppressing cell proliferation. This incongruity can be explained by two theories on the cAMP signalling cascade. The first theory proposes that elevation of intracellular cAMP is beneficial for suppressing cell proliferation in most mesenchymal and epithelial cell lines, such as glioblastoma (Kang et al. 2014), thyroid cells (Sawa et al. 2017), lung and breast carcinoma cells (Lerner and Epstein 2006), ovarian granulosa cells (Zwain and Amato 2001), fibroblasts (Huston et al. 2006), and primary cardiomyocytes (Ding et al. 2005). In contrast, the second theory proposes that cAMP promotes cell survival, which has been observed in myeloid cells, pancreatic β-cells, hepatocytes, gastric and intestinal cells, spinal motor, superior cervical ganglion sympathetic, dorsal root ganglion, dopaminergic neurons, cerebral granule and septal cholinergic neurons (Kwon et al. 2004, Lerner and Epstein 2006, Cullen et al. 2004, Nishihara et al. 2003, Hoshino et al. 2003, Chin and D’Mello 2005). These two divergent roles of cAMP may be crucial in both physiological maintenance and pathological conditions, but whether these signalling cascades are interconnected remains unclear.

It has been well established that after synthesis by AC activation, cAMP diffuses within cells and is hydrolyzed into inactive forms by phosphodiesterases (PDEs). PDEs are a subfamily of ectonucleotidases consisting of 11 mammalian isoforms (PDE1–11) in mammals and are encoded by 21 different genes (Zhang 2015, Beavo 1995), which are distributed in many types of tissue (Torphy 1998, Soderling and Beavo 2000, Mehats et al. 2002, Houslay and Adams 2003, Castro et al. 2005, Goraya and Cooper 2005, Ke and Wang 2007). Each isoenzyme possesses different affinities for cAMP and/or cGMP and kinetic characteristics that result in their distinctive response to a stimulus (Menniti et al. 2006). To date, there are 3 classes of PDEs subdivided according to their substrate specificities: cAMP-specific PDEs (PDE4, PDE7 and PDE8), cGMP-specific PDEs (PDE5, PDE6 and PDE9), and dual-substrate PDEs (PDE1, PDE2, PDE3, PDE10 and PDE11) (Francis et al. 2011). Through metabolizing both cAMP and cGMP, PDEs generate intracellular gradients and microdomains of these second messengers to regulate their spatio-temporal signalling (Menniti et al. 2006, Conti et al. 2014). PDEs prevent non-specific activation enabling both specificity and selectivity towards intracellular targets (Ladilov and Appukuttan 2014).

Altered pattern on cAMP in the brain has been reported by overexpression of some PDEs, such as PDE1, PDE4, PDE5, and PDE7 (Sengupta et al. 2011, Cesarini et al. 2017, Savai et al. 2010, Vatter et al. 2005, Brooks et al. 2014). Some evidence shows that particular PDE inhibitors, such as rolipram, a selective PDE4 inhibitor, prevents leukaemia proliferation through an elevation of cAMP and an induction of apoptosis. This suggests that using specific PDE inhibitors is a viable approach for cancer therapy (Kuida et al. 1996, Chen et al. 2007). Given that PDE inhibitors may offer therapeutic efficacy against cancer, we investigated the role of each PDE upon cAMP accumulation and cell proliferation in glioma cell models using a range of pharmacological inhibitors. Our data indicates that tumour cell regression is linearly correlated with elevation of cAMP. Among all PDE inhibitors tested, trequinsin (a non-selective PDE2/3/7 inhibitor (Rickles et al. 2010)) was found to be the most potent at inhibiting cell proliferation. More importantly, by using small molecule compounds we highlight that simultaneous elevation of cAMP through activation of AC and inhibition of multiple PDEs (specifically PDE2, PDE3 and PDE7) had synergistic anti-proliferative effects on the glioma cells, predominantly by altering cell cycle progression and inducing activation of caspase-3/7, providing a novel treatment strategy for glioma.

## 2. METHODS

### Cell Lines

C6 glioma cells (a gift from Prof. Colin Taylor, University of Cambridge) were cultured in Gibco® MEM (Thermo Fisher, UK) supplemented with 10% foetal bovine serum (FBS, Sigma, UK), 2 mM L-glutamine (Sigma, UK), and 1% antibiotic/antimycotic (Sigma, UK). ST14A cells (rat-derived striatal cells (Tissue and Cell Biotechnologies, Italy) were grown in Gibco® DMEM/F12 1:1 (1X) – Glutamax TM (Thermo Fisher, UK), supplemented with 10% FBS and 1% antibiotic/antimycotic. C6 cells were maintained at 37 °C in humidified 95% air and 5% CO_2_. ST14A cells were grown at 33 °C in humidified 95% air and 5% CO_2_ because propagation of ST14A cells at 37 °C has been shown to induce differentiation into glial cells (Ehrlich et al. 2001). Where appropriate cells were treated with pertussis toxin (PTX, Thermo Fisher, UK) at a range concentration of 2 pg/ml to 200 ng/ml or cholera toxin (CTX, Sigma, UK), at a concentration of 3.5 pg/ml to 350 ng/ml. PTX uncouples receptor-mediated Gα_i_-dependent inhibition of cAMP production (Weston et al. 2016). CTX inhibits GTPase activity of Gα_s_ and causes permanent activation.

### Compounds

Forskolin (Sigma, U.K.) was diluted to a stock of 100 mM in DMSO (Sigma, U.K.). All PDE inhibitors were initially supplied by IOTA Pharmaceuticals. The names of the compounds were only known to IOTA Pharmaceuticals and initial experiments (Figures 2 and 3) were performed blind. Upon data analysis, IOTA Pharmaceuticals decoded the compounds to enable subsequent characterisation. Trequinsin, PF-2545920, vinpocetine, sildenafil, rolipram, cilostamide, caffeine, SQ22536, EHNA, amrinone, zaprinast, TC3.6, ibudilast, milrinone, BAY 73-6691, BRL-50481, piclamilast, IBMX, roflumilast, tadalafil, PF-04449613 (all purchased from Sigma, UK), BC 11-38, and PF 04671536 (both obtained from Tocris, UK) were dissolved in DMSO to stock concentrations of either 100 mM or 10 mM. Guanylyl cyclase activators BAY 41-8543 and YC-1 were purchased from Tocris and diluted to a stock of 100 mM in DMSO. Epac inhibitors ESI-09, HJC0350, and CE3F4 (all purchased from Sigma, UK) were diluted to 10 mM stocks in DMSO. PKA and PKG inhibitors, KT5720 and KT5823, respectively (all obtained from Cambridge Insight Biotechnology, UK) were diluted to stocks of 10 mM and 1 mM, respectively.

**Figure 1.**
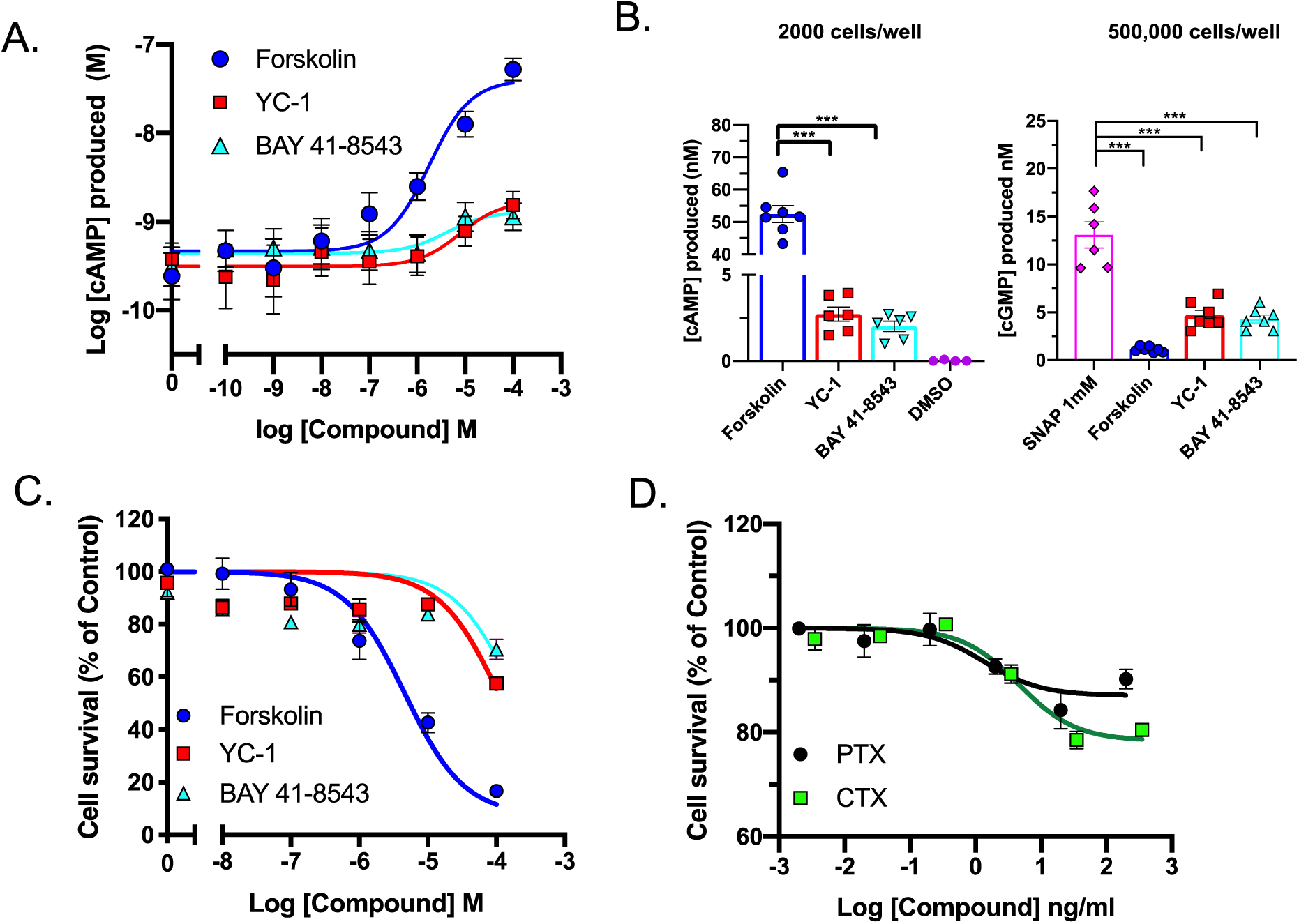
Elevation of cAMP, but not cGMP, mediates cell growth suppression. *A*. Representative cAMP dose-response curves following 30 minutes treatment with adenylyl cyclase activator (forskolin) or guanylyl cyclase activators (YC-1 and BAY 41-8543). *B*. Comparison of accumulation of cAMP and cGMP in C6 cells in response to forskolin (100 µM), BAY 41-8543 (100 µM), YC-1 (100 µM), or SNAP (100 µM). Cell survival following 72 hours treatment with forskolin, BAY 41-8543 or YC-1 (C) or PTX or CTX (D). Data are expressed as percentage survival relative to vehicle alone (mean±SEM) of 6-9 individual experiments. Statistical significance of differences from the parameters measured in the presence of either forskolin or SNAP was assessed by the one-way analysis of variance followed by the Dunnett’s *post hoc* test (***, p<0.001).

**Figure 2.**
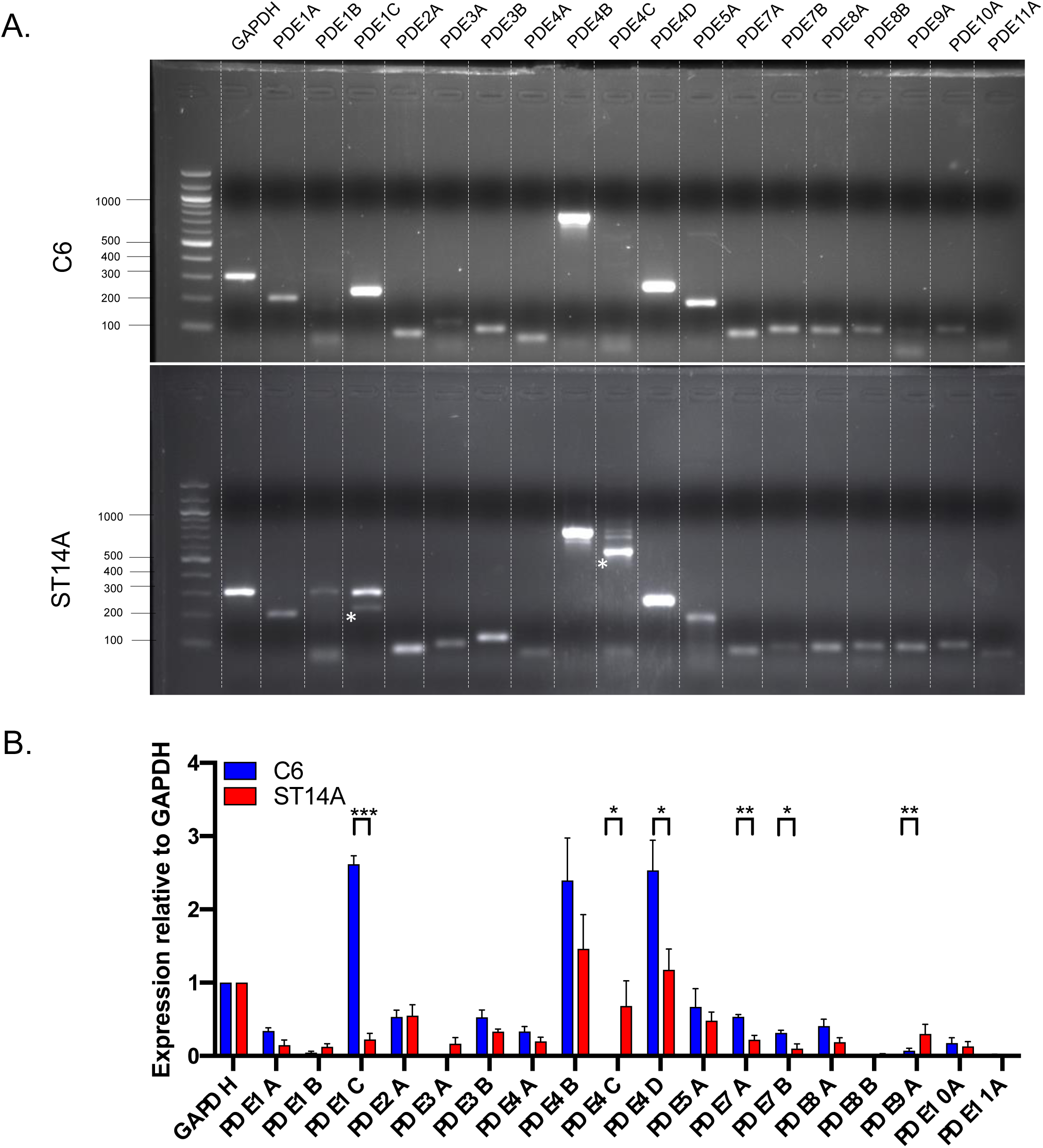
Expression profile of PDE isoenzymes in C6 and ST14A cells. *A.* Representative gel documentation showing amplified PDEs genes from C6 and ST14A cell line. (*) on the gel showed correct band size which have been confirmed by sequencing. *B.* Semi-quantitative mRNA levels in C6 cells and ST14A cells. Expression of each gene of interest was normalised relative to GAPDH. Data are expressed as the mean ± SEM from 5-7 individual repeats. Data were determined as statistically different (*, p<0.05; **, p<0.01; ***, p<0.001) compared to individual isoenzyme between both cell lines using Student’s t-test analysis.

**Figure 3.**
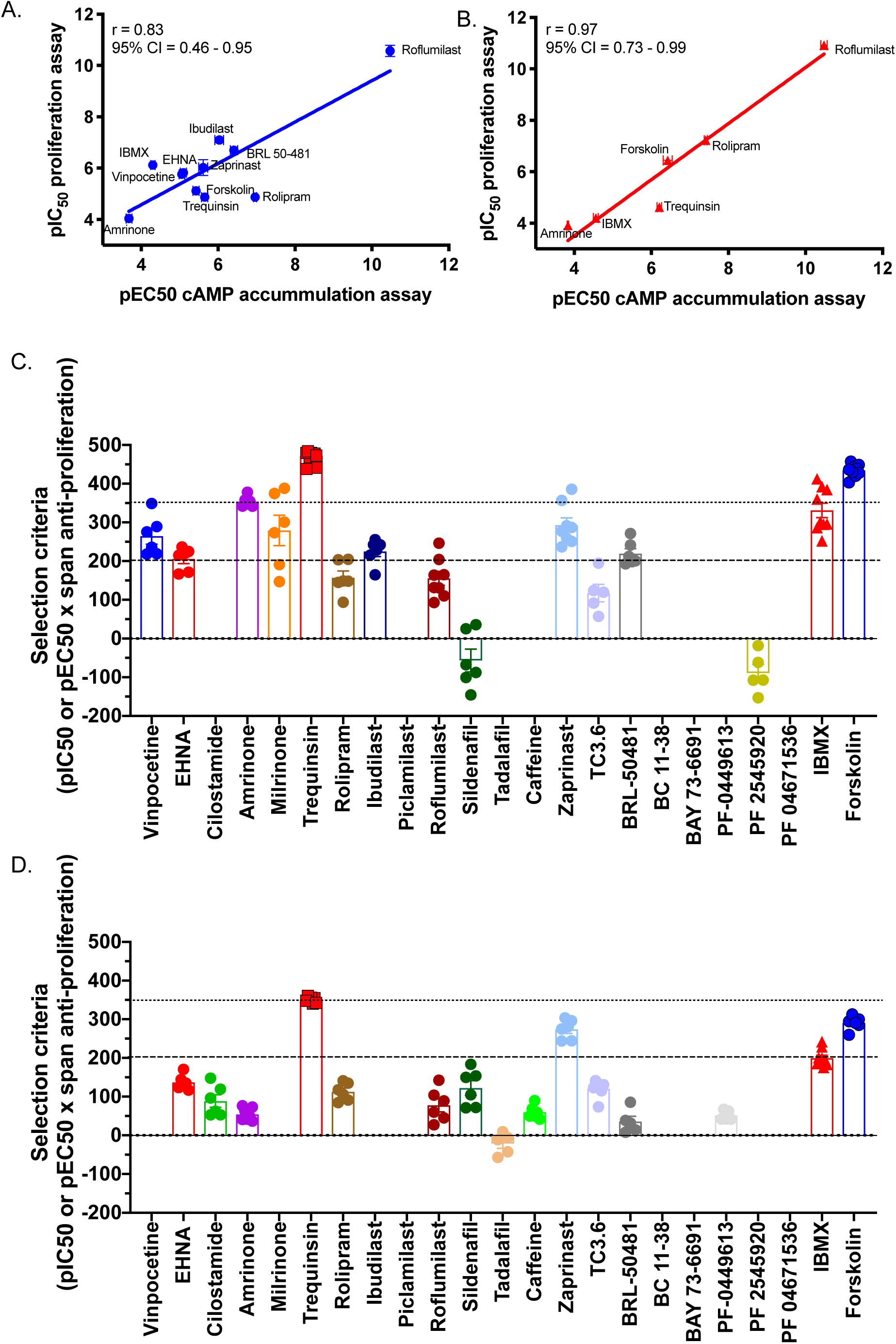
Elevation of intracellular cAMP is positively correlated with cell growth suppression. *A-B.* Correlation (with 95% confidence interval) of log potencies of each PDE inhibitor in C6 (A) and ST14A cells (B) was determined by calculating Pearson’s correlation coefficient (r). *C-D.* Compound selection criteria from C6 (C) and ST14A cells (D) was calculated based on potency and efficacy in proliferation assay. The dashed lines represent threshold value of 200 (less stringent criteria) and the dotted lines a higher criteria value of 350. Individual data point was obtained from supplemental information (Figure 1-4).

### Reverse Transcription PCR

RNA was extracted from C6 and ST14A cells using RNAqueous®-4PCR Total RNA Isolation Kit (Life Technologies, UK) according to manufacturer’s instructions. In order to remove any contamination of genomic DNA, all RNA samples were treated with DNAse I included in the kit. The purity of RNA samples was quantified using a NanoDrop ™Lite spectrophotometer (Thermo Scientific, UK) and only samples that had a minimum yield of 100 ng/μL and A_260/280_ >1.9 were used in the experiments. Complementary DNA was synthesized using a QuantiTect reverse transcription kit (Qiagen, UK). The oligonucleotides (Sigma, UK) used for PCR were designed specifically for rat and the sequences are displayed at methods section.

The oligonucleotides (Sigma, UK) used for PCR were designed specifically for rat and the sequences are as follows: GAPDH: forward 5′-TCCCTCAAGATTGTCAGCAA-3′, reverse 5′- AGATCCACAACGGATACATT-3′ (309 bp); PDE1A: forward 5′- CGCCTGAAAGGAATACTAAGA-3′, reverse 5′-TAGAAGCCAACCAGTCCCGGA-3′ (211 bp); PDE1B: forward 5′- CTGTCACCCCGCAGTCCTCCG-3′, reverse 5′- GAAGGTGGAGGCCAGCCAGTC-3′ (309 bp); PDE1C forward 5′- CGCGGGCTGAGGAAATATAAG-3′, reverse 5′-GAAGGTGGAGGCCAGCCAGTC-3′ (237 bp); PDE2A: forward 5′-CCAAATCAGGGACCTCATATTCC-3′, reverse 5′- GGTGTCCCACAAGTTCACCAT-3′ (86 bp); PDE3A: forward 5′- CACAAGCCCAGAGTGAACC-3′, reverse 5′-TGGAGGCAAACTTCTTCTCAG-3′ (123 bp); PDE3B: forward 5′-GTCGTTGCCTTGTATTTCTCG-3′, reverse 5′- AACTCCATTTCCACCTCCAGA-3′ (103 bp); PDE4A: forward 5′- CGACAAGCACACAGCCTCT-3′, reverse 5′-CTCCCACAATGGATGAACAAT-3′ (73 bp); PDE4B: forward 5′-CAGCTCATGACCCAGATAAGTGG-3′, reverse 5′- GTCTGCACAAGTGTACCATGTTGCG-3′ (787 bp); PDE4C: forward 5′- ATGGCCCAGATCACTGGGCTGCGG-3′, reverse 5′- GCTGAGGTTCTGGAAGATGTCGCAG-3′ (582 bp); PDE4D: forward 5′- CCCTCTTGACTGTTATCATGCACACC-3′, reverse 5′- GATCCTACATCATGTATTGCACTGGC-3′ (262 bp); PDE5A: forward 5′- CCCTGGCCTATTCAACAACGG-3′, reverse 5′-ACGTGGGTCAGGGCCTCATA-3′ (192 bp); PDE7A: forward 5′-GAAGAGGTTCCCACCCGTA-3′, reverse 5′- CTGATGTTTCTGGCGGAGA-3′ (85 bp); PDE7B: forward 5′-GGCTCCTTGCTCATTTGC-3′, 5′-GGAACTCATTCTGTCTGTTGATG-3′ (99 bp); PDE8A: forward 5′- TGGCAGCAATAAGGTTGAGA-3′, reverse 5′-CGAATGTTTCCTCCTGTCTTT-3′ (97 bp); PDE8B: forward 5′-CTCGGTCCTTCCTCTTCTCC-3′, 5′-AACTTCCCCGTGTTCTATTTGA-3′ (147 bp); PDE9A: forward 5′-GTGGGTGGACTGTTTACTGGA-3′, reverse 5′- TCGCTTTGGTCACTTTGTCTC-3′ (107 bp); PDE10A: forward 5′- GACTTGATTGGCATCCTTGAA-3′, reverse 5′-CCTGGTGTATTGCTACGGAAG-3′ (115 bp); and PDE11A: forward 5′-CCCAGGCGATAAATAAGGTTC-3′, reverse 5′- TGCCACAGAATGGAAGATACA-3′ (87 bp).

All PCR products were run on a 2% agarose gel. The gel was visualised in the presence of ethidium bromide and imaged using a G Box iChemi gel documentation system. Density of each band was analysed with GeneTools analysis software (Syngene, UK). In order to confirm the gene of interest was correctly amplified, gel extraction was performed using Qiaquick Gel Extraction kit (Qiagen, UK) and the samples sequenced to confirm fidelity (inhouse sequencing service).

### cAMP accumulation assay

Confluent cells were trypsinised and re-suspended in stimulation buffer (PBS with 0.1% BSA). Cells were plated onto 384-well optiplates (Perkin Elmers, UK) at a density of 2000 cells/well. To determine the efficacy of individual PDE inhibitor, cells were co-stimulated with three different concentrations of compounds (which spanned 100-fold either side of the individual IC_50_ value in vitro) and pEC_20_ values of forskolin (1.6 μM for C6 cells and 50 nM for ST14A cells) for 30 minutes. Stimulating cells with the pEC_20_ of forskolin, enables a larger range to observe any effect of the PDE inhibitors on cAMP production. To generate full dose response curves, compounds were added to cells in the range of 0.1 pM – 100 µM for 30 minutes. Detection of cAMP was assessed using LANCE cAMP detection kit and end-point measurement was performed using a Mithras LB940 microplate reader (Berthold Technologies, Germany). The lysed cells were excited at 340 nm wavelength with fluorescence from homogeneous time-resolved FRET detected at 665 nm. All data were normalised to the maximal level of cAMP accumulation from cells in response to 100 μM forskolin stimulation and some were also interpolated to cAMP standard curve.

### cGMP accumulation assay

Confluent C6 cells were trypsinised and resuspended in PBS containing 0.1% BSA. Cells were plated onto a 384-well plate at a density of 500,000 cells/well and stimulated with compounds for 30 minutes. After stimulation, 5 µl of d2-cGMP analogue and 5 µl mAb-cryptate were added to each well and incubated for 1 hour at room temperature as per the manufacturer’s instruction (Cisbio, France). The d2-cGMP fluorophore was excited at a wavelength of 337 nm and emission was detected at 665 nm and 620 nm. Fluorescence was measured using a Mithras LB940 microplate reader (Berthold Technologies, Germany). Delta F% values were calculated using the 665 nm/620 nm ratio and all data were interpolated to a standard curve which covered an average cGMP range of 0.5 – 50 nM.

### Cell proliferation assay

C6 glioma cells were seeded at a density of 2,500 cells/well in a clear flat bottom 96-well plate (Corning). After 24 hours, cells were exposed to test compounds or vehicle, as indicated, and were incubated for 72 hours. To further investigate whether downstream pathways of cAMP influenced cell proliferation, cells were cotreated with selective inhibitors that target cAMP/cGMP sensors including PKA, PKG, and Epac. Cells were seeded as previously described and treated with either forskolin or trequinsin in the presence of the following inhibitors: KT5720 to inhibit PKA, KT5823 to inhibit PKG, ESI-09 as non-selective Epac inhibitor, CE3F4 as a selective Epac1 inhibitor, and HJC0350 as a selective Epac2 inhibitor.

After 72 hours incubation, 5 μL of Cell Counting Kit – 8 (CCK-8, Sigma, UK) was added to each well and the cells were incubated for an additional 2-3 hours at 37 °C in the dark. The absorbance of each well was measured using a Mithras LB940 microplate reader (Berthold Technologies, Germany) with an excitation of 450 nm. The amount of formazan formed is directly proportional to the number of viable cells. Cell proliferation was calculated as a percentage of number of cells treated with vehicle alone.

### Caspase assay

Cells were seeded in to clear bottom black 96-well plates (Corning) and treated with forskolin (1-100 μM), trequinsin (1–100 μM) or staurosporine (1 μM, a pan caspase activator) in complete MEM media. 1% DMSO was used as vehicle control. Cells were exposed to test compounds for 72 hours, plates were treated with 2 μM of the CellEvent™Caspase-3/7 green detection reagent (Life Technologies, UK) for 60 minutes at 37 °C in the dark. Caspase activity was detected by cleavage of the tetrapeptide substrate DEVD, which is conjugated to a nucleic acid binding dye. Intracellular caspase-3/7 activities were imaged using a BD Pathway 855. To normalise the number of cells with caspase activated, cells were also labelled with Hoechst 33342 (Cambridge Bioscience, UK). Activated caspase-3/7 cleaves substrate and produce green fluorescence which was visualised using FITC/Alexa Fluor™488 filter setting. The total number of cells stained with Hoechst 33342 was measured using Hoechst filter (350/461 nM).

### Cell Cycle Analysis

Cell cycle analysis using flow cytometry provides information on the distribution of cells in interphase stages of the cell cycle (G_0_/G_1_, S, and G_2_/M). C6 cells were seeded in to 24-well plates and cultured for 24 hours. Cells were exposed to selected treatments including forskolin, trequinsin, and a combination of individual PDE2,3,7 inhibitor, for 72 hours. Subsequently, cells were harvested and resuspended in PBS containing 0.1% Triton X-100, 10 µg/ml RNase A, and 5 µg/ml propidium iodide (PI) before incubation at 37 °C for 15 minutes. Samples were analysed using a BD Accuri C6 flow cytometer and cell cycle analysis was performed using BD C6 software.

### Statistical analysis

To quantify gene expression through RT-PCR, the densitometry results of each gene of interest were normalised to GAPDH signal. For cAMP accumulation and cell proliferation assays, normalised data were fitted to obtain concentration–response curves using the three-parameter logistic equation using GraphPad Prism 8 (Graphpad Software, San Diego) to obtain values of E_max_ /I_max_, pEC_50_/pIC_50_, baseline, and span. Statistical differences were analysed using one-way ANOVA followed by Dunnett’s *post-hoc* (for comparisons amongst more than two groups) or independent Student’s t-test (for comparison between two groups). To determine the correlation of cAMP levels and cell proliferation of each PDE inhibitor in both C6 and ST14A cells, Pearson’s correlation coefficient (r) was calculated with 95% confidence interval.

## 3. RESULTS

### 3.1 Elevation of intracellular cAMP levels reduces cell proliferation in a glioma cell line

We first sought to determine if changes in the intracellular cAMP concentration modulated glioma cell growth. When C6 cells were exposed to forskolin, we observed dose-dependently increases of intracellular cAMP (Figure 1A, B) and reduced cell proliferation (Figure 1C).

Given the fact that there is cross-talk between the cAMP and cGMP pathways, we also evaluated the role of the cGMP pathway on cell proliferation by treating cells with the small molecule guanylyl cyclase activators, BAY 41-8543 and YC-1. Both compounds elevated intracellular cGMP levels (Figure 1B), however, the cGMP production was 1000x lower than cAMP production in C6 cells even in response to treatment with the nitric oxide (NO) donor, S-Nitroso-N-acetyl-DL-penicillamine (SNAP). Surprisingly, both BAY 41-8543 and YC-1 also dose-dependently increased cAMP levels (Figure 1A) but did not exceed ∼20% response relative to that of forskolin (Figure 1A). BAY 41-8543 and YC-1 also had a minimal effect on cell proliferation compared to forskolin, with a reduction in cell survival only observed at 100 µM (Figure 1C). These anti-proliferative effects may occur due to modulatory effect between cAMP and cGMP. Accumulation of cGMP levels are possible to be a regulator of the activity of dual-substrate PDEs leading to potentiation of cAMP, thereby cell growth suppression was observed. These results suggest that that elevation cAMP pathway plays a more important role in reducing cell proliferation.

Heterotrimeric G proteins are the primary effectors of G protein-coupled receptors (GPCRs). Activation of Gα_s_ is well known to activate AC to stimulate intracellular cAMP production, whilst activation of Gα _i/o_ inhibits AC and cAMP synthesis (Gilman 1990, Hildebrandt 1997). Pertussis toxin (PTX) and cholera toxin (CTX) were utilised to determine whether G protein-mediated cAMP production inhibits cell growth in C6 cells. Cell survival of C6 cells was only suppressed by 20% after treatment with PTX or CTX in a dose-dependent manner (Figure 1D). Together, these results suggest that increasing intracellular cAMP concentration, through direct activation of AC, plays a pivotal role in inhibiting cell growth with a minimal involvement of cGMP.

### 3.2 Specific inhibition of PDEs indicates reliance on PDE expression levels

Given that intracellular concentrations of cAMP are modulate both by its production and its degradation, we next sought to investigate the expression of PDEs in glioma cell line. We performed reverse transcription PCR (rt-PCR) to determine the expression of each PDE isoenzyme in C6 cells, as a model for glioma derived from rat compared to ST14A cells, as a rodent model for healthy neurons. As shown in Figure 2, the overall expression level of PDEs was higher in C6 cells compared to ST14A cells. With the exception of PDE1C there was little overall difference in the profiles of PDEs expressed between C6 cells and ST14A cells. PDE1C, PDE4D, PDE7A, and PDE7B were the only PDEs to display significant elevation in the C6 cells.

The rt-PCR expression profiles showed a wide number of PDEs to be expressed in the glioma cells. However, this was not quantitative thereby not providing a clear indication as to which ones might be the most important. Thus, we next, investigated the role of PDEs in regulating cAMP levels and cell proliferation in C6 cells by applying small molecule selective PDE inhibitors as tools to modify their action (Figure 3A–D and supplementary Figures). The small compounds were screened blindly and subsequently decoded after data analysis (see methods). Both cell lines were stimulated with the pEC_20_ concentration of forskolin. Stimulating cells with a low concentration of a pan-AC activator increases the possibility of detecting an elevation of cAMP in the presence of selected PDE inhibitor. The pharmacological parameters of each PDE inhibitor could then be determined using the three parameters non-linear regression equation to fit full dose-response curves using Prism 8.0. pEC_50_ values for cAMP production and pIC_50_ values for cell growth inhibition for each compound are quoted in Table 1 and their pharmacological actions are summarised in Table 2.

**Table 1.** Potency values for cAMP production (pEC_50_) and for cell growth inhibition (pIC_50_) of each PDE inhibitor in C6 glioma cells and ST14A cells.

| Compounds | PDE Target | C6 |  |  |  | ST14A |  |  |  |
| --- | --- | --- | --- | --- | --- | --- | --- | --- | --- |
|  |  | pEC<sub>50</sub> | Span (cAMP) | pIC<sub>50</sub> | Span (proliferation) | pEC<sub>50</sub> | Span (cAMP) | pIC<sub>50</sub> | Span (proliferation) |
| Vinpocetine | 1 | 5.10 ± 0.08* | 27.86 ± 1.05*** | 5.83 ± 0.13* | 45.33 ± 3.12*** | 4.16 ± 0.04*** | 13.15 ± 2.22*** | N/A | N/A |
| EHNA | 2 | 5.06 ± 0.07* | 24.26 ± 1.78*** | 5.76 ± 0.22 | 36.34 ± 3.31*** | N/A | N/A | 6.57 ± 0.08*** | 20.86 ± 1.20*** |
| Cilostamide | 3 | 7.19 ± 0.09*** | 15.20 ± 1.45*** | N/A | N/A | N/A | N/A | 8.39 ± 0.15*** | 10.56 ± 1.90 |
| Amrinone | 3 | 3.68 ± 0.08*** | 108.9 ± 4.82*** | 4.04 ± 0.08*** | 86.44 ± 2.26 | 3.84 ± 0.02*** | 89.70 ± 1.12** | 3.91 ± 0.11*** | 54.37 ± 6.17*** |
| Milrinone | 3 | N/A | N/A | 8.06 ± 0.13*** | 34.49 ± 4.66*** | 7.05 ± 0.18*** | 27.45 ± 2.35 | N/A | N/A |
| Trequinsin | 2,3,7 | 5.65 ± 0.08 | 50.57 ± 2.02*** | 4.87 ± 0.07 | 96.93 ± 0.94 | 6.21 ± 0.06 | 80.31 ± 1.85 | 4.62 ± 0.04*** | 75.82 ± 0.79*** |
| Rolipram | 4 | 6.96 ± 0.04*** | 60.46 ± 3.74 | 6.78 ± 0.20*** | 23.44 ± 2.94*** | 7.41 ± 0.04*** | 59.26 ± 4.11** | 7.23 ± 0.05*** | 15.57 ± 1.28*** |
| Ibudilast | 4 | 6.03 ± 0.11*** | 64.21 ± 3.46 | 7.05 ± 0.17*** | 31.76 ± 1.30*** | 6.15 ± 0.12 | 48.64 ± 2.18*** | N/A | N/A |
| Piclamilast | 4 | 8.72 ± 0.04*** | 41.32 ± 2.87*** | N/A | N/A | 9.02 ± 0.14*** | 35.24 ± 1.96*** | N/A | N/A |
| Roflumilast | 4 | 10.47 ± 0.03*** | 54.58 ± 3.12 | 10.57 ± 0.22*** | 14.72 ± 1.66*** | 10.48 ± 0.08*** | 28.77 ± 0.78*** | 10.92 ± 0.09*** | 7.19 ± 1.63*** |
| Sildenafil | 5 | N/A | N/A | 8.56 ± 0.16*** | -6.77 ± 3.54***# | 8.79 ± 0.20*** | -4.57 ± 0.93***# | 9.67 ± 0.11*** | 12.71 ± 2.02*** |
| Tadalafil | 5 | N/A | N/A | N/A | N/A | 7.88 ± 0.04*** | -23.15 ± 1.38***# | 8.81 ± 0.43*** | -2.69 ± 1.55***# |
| Caffeine | 1,4,5 | N/A | N/A | N/A | N/A | N/A | N/A | 6.65 ± 0.06*** | 9.02 ± 1.08*** |
| Zaprinast | 5,6,9,11 | 5.61 ± 0.11 | 14.27 ± 1.12*** | 6.03 ± 0.30*** | 49.70 ± 4.60*** | N/A | N/A | 5.86 ± 0.09*** | 46.71 ± 1.45*** |
| TC3.6 | 7 | N/A | N/A | 7.53 ± 0.26*** | 15.31 ± 2.47*** | N/A | N/A | 5.69 ± 0.05*** | 21.22 ± 1.49*** |
| BRL-50481 | 7 | 6.41 ± 0.09*** | 31.3 ± 1.67*** | 6.70 ± 0.15*** | 33.07 ± 2.74*** | N/A | N/A | 7.73 ± 0.24*** | 4.52 ± 1.68*** |
| BC 11-38 | 8 | N/A | N/A | N/A | N/A | N/A | N/A | N/A | N/A |
| BAY-736691 | 9 | N/A | N/A | N/A | N/A | N/A | N/A | N/A | N/A |
| PF-0449613 | 9 | N/A | N/A | N/A | N/A | 7.55 ± 0.12*** | -23.48 ± 1.75***# | 8.05 ± 0.08*** | 6.37 ± 0.72 |
| PF 2545920 | 10A | N/A | N/A | 9.46 ± 0.06*** | -8.97 ± 3.05# | N/A | N/A | 10.78 ± 0.11*** | 11.94 ± 4.13 |
| PF 04671536 | 11 | N/A | N/A | N/A | N/A | N/A | N/A | N/A | N/A |
| IBMX | Non-selective | 4.30 ± 0.06*** | 42.79 ± 1.53*** | 6.12 ± 0.06*** | 54.17 ± 3.20 | 4.56 ± 0.06*** | 57.53 ± 3.52 | 4.20 ± 0.09 | 47.95 ± 2.70 |
| Forskolin | AC activator | 5.43 ± 0.08 | 64.97 ± 3.51 | 5.12 ± 0.06 | 84.70 ± 1.44 | 6.43 ± 0.12 | 74.06 ± 3.07 | 6.44 ± 0.02 | 45.11 ± 1.22 |

N/A – not applicable; compounds did not have any effect on cAMP production or cell growth inhibition. Data are presented as the mean ± SEM of 6-9 individual repeat. Data were determined as statistically different (\*, p<0.05; \*\*\*, p<0.001) compared to forskolin using one-way ANOVA followed by Dunnett's *post-hoc* analysis. #, showing negative responses, either suppressing cAMP accumulation or being proliferative

**Table 2.** Summary of the pharmacological effects of each PDE inhibitor on cAMP production and cell proliferation.

| Compound | Target | C6 |  | ST14A |  |
| --- | --- | --- | --- | --- | --- |
|  |  | cAMP | Proliferation | cAMP | Proliferation |
| Vinpocetine | 1 | ↑ | ↓↓ | ↑ | - |
| EHNA | 2 | ↑ | ↓↓ | ↓ | ↓ |
| Cilostamide | 3 | ↑ | - | ↓ | ↓ |
| Amrinone | 3 | ↑↑↑ | ↓↓↓ | ↑↑↑ | ↓↓↓ |
| Milrinone | 3 | - | ↓↓ | ↑ | - |
| Trequinsin | 2,3,7 | ↑↑↑ | ↓↓↓ | ↑↑↑ | ↓↓↓ |
| Rolipram | 4 | ↑↑↑ | ↓ | ↑↑↑ | ↓ |
| Ibudilast | 4 | ↑↑↑ | ↓↓ | ↑↑ | - |
| Piclamilast | 4 | ↑↑ | - | ↑↑ | - |
| Roflumilast | 4 | ↑↑↑ | ↓ | ↑ | ↓ |
| Sildenafil | 5 | - | ↑ | ↓ | ↓ |
| Tadalafil | 5 | - | ↑ | ↓ | ↑ |
| Caffeine | 1,4,5 | Bell-shape | - | - | ↓ |
| Zaprinast | 5,6,9,11 | ↑ | ↓↓ | - | ↓↓ |
| TC3.6 | 7 | - | ↓ | - | ↓ |
| BRL-50481 | 7 | ↑↑ | ↓↓ | - | ↓ |
| BC 11-38 | 8 | - | - | - | - |
| BAY-736691 | 9 | - | - | - | - |
| PF-0449613 | 9 | - | - | ↓ | ↓ |
| PF 2545920 | 10A | - | ↑ | - | ↓ |
| PF 04671536 | 11 | - | - | - | - |
| IBMX | Non-selective | ↑↑ | ↓↓↓ | ↑ | ↓↓ |

↑ = increase 10-30%, ↑↑ = increase 31-50%, ↑↑↑ = increase >50%, ↓: suppress 10-30%, ↓↓: suppress 31-50%, ↓↓↓: suppress >50%

Plotting the potencies of each PDE inhibitor revealed a significant (p<0.001) positive correlation between elevation of intracellular cAMP and inhibition of cell growth (Figure 3A and B) for both C6 cells (r = 0.83 (95% confidence interval 0.46–0.95)) and ST14A cells (0.97 (95% confidence interval 0.73–0.99)). To provide a convenient and rapid method for comparing the each PDE inhibitor we used a simple algorithm whereby we combined (multiplied) the terms for efficacy and affinity in proliferation assay by using potency and span values. An ideal compound would be one that shows high potent for inhibition of growth and a large range. Using the data in table 1 we calculated these values (arbitrary units – Figure 3C and 3D) for both C6 and ST14A cells. We set an arbitrary threshold of 200 to determine compounds that might be worth further investigation. From our initial screen only 9 compounds were deemed to have passed the threshold vinpocetine (PDE1 inhibitor), amrinone, milrinone (both PDE3 inhibitors), ibudilast (PDE4 inhibitor), trequinsin, IBMX and zaprinast (multiple PDE isoform inhibitors), BRL73-6691 (PDE7 inhibitor), and foskolin (pan-AC activator). Comparison with expression data C6 cells (Figure 2A and 2B) shows good correlation with the inhibitors that were successful in the screen. It is worth noting that the compounds that showed the highest values (>350) were forskolin, trequinsin and IBMX. These compounds are suggested to target multiple components in the cAMP synthesis/degradation pathway so explaining their great affinity/efficacy. From these data two compounds were discontinued IBMX and zaprinast due to selectivity issue. IBMX has been reported to be a ligand to adenosine receptor (Morgan et al. 1993), whereas zaprinast is also known to bind to GPR35 (Taniguchi et al. 2006). Finally, it is worth noting that all compounds displayed lower affinity/efficacy values in ST14a cells (Figure 3D), and this is consistent with the reduced PDE expression compared to C6 cells previously described (Figure 2B).

### 3.3 Inhibitor cocktail of individual PDE2, PDE3, and PDE7 inhibitor exhibited similar effect to that of trequinsin, both upon elevating cAMP levels and suppressing cell growth

Of all the compounds tested, our data shows that trequinsin is the most potent compound at increasing intracellular cAMP levels and suppressing cell growth. Trequinsin is known to potently inhibit PDE3, but it may also block the cAMP binding site of PDE2 and PDE7 (Rickles et al. 2010). This, however, has not been thoroughly investigated. Therefore, we probed the mechanism by which trequinsin exerts its antiproliferative effects by combining selective inhibitors against PDE2, PDE3, and PDE7 – by using EHNA, amrinone, and BRL-50481, respectively.

None of the selective PDE inhibitors were more potent at stimulating cAMP accumulation or inhibiting cell proliferation, as individual treatments, than trequinsin in C6 cells (Figure 4A and 4D). Although amrinone displayed a similar E_max_ to that of trequinsin for cAMP accumulation, the I_max_ for cell growth inhibition was ∼50% reduced relative to trequinsin (Figure 4A and 4D). Subsequently, we investigated the combinatorial effect of individual PDE2, 3, and 7 inhibitors. The combination of EHNA and amrinone (PDE2 and PDE3 inhibitors), as well as EHNA and BRL-50481 (PDE2 and PDE7 inhibitors), enhanced the potency of effect compared to when these drugs were used individually (Figure 4B and 4E). However, the combination of BRL-50491 with amrinone (PDE3 and PDE7 inhibitors) was comparable to that of amrinone alone. Interestingly, when all three selective inhibitors were combined the potency and efficacy were similar to that of trequinsin (Figure 4C and 4F). Overall, this suggests that trequinsin induces its antiproliferative effects through the simultaneous inhibition of PDE2, PDE3 and PDE7. It is trequinsin’s multiple PDE isoform inhibition that provides its superior activity in C6 cells.

**Figure 4.**
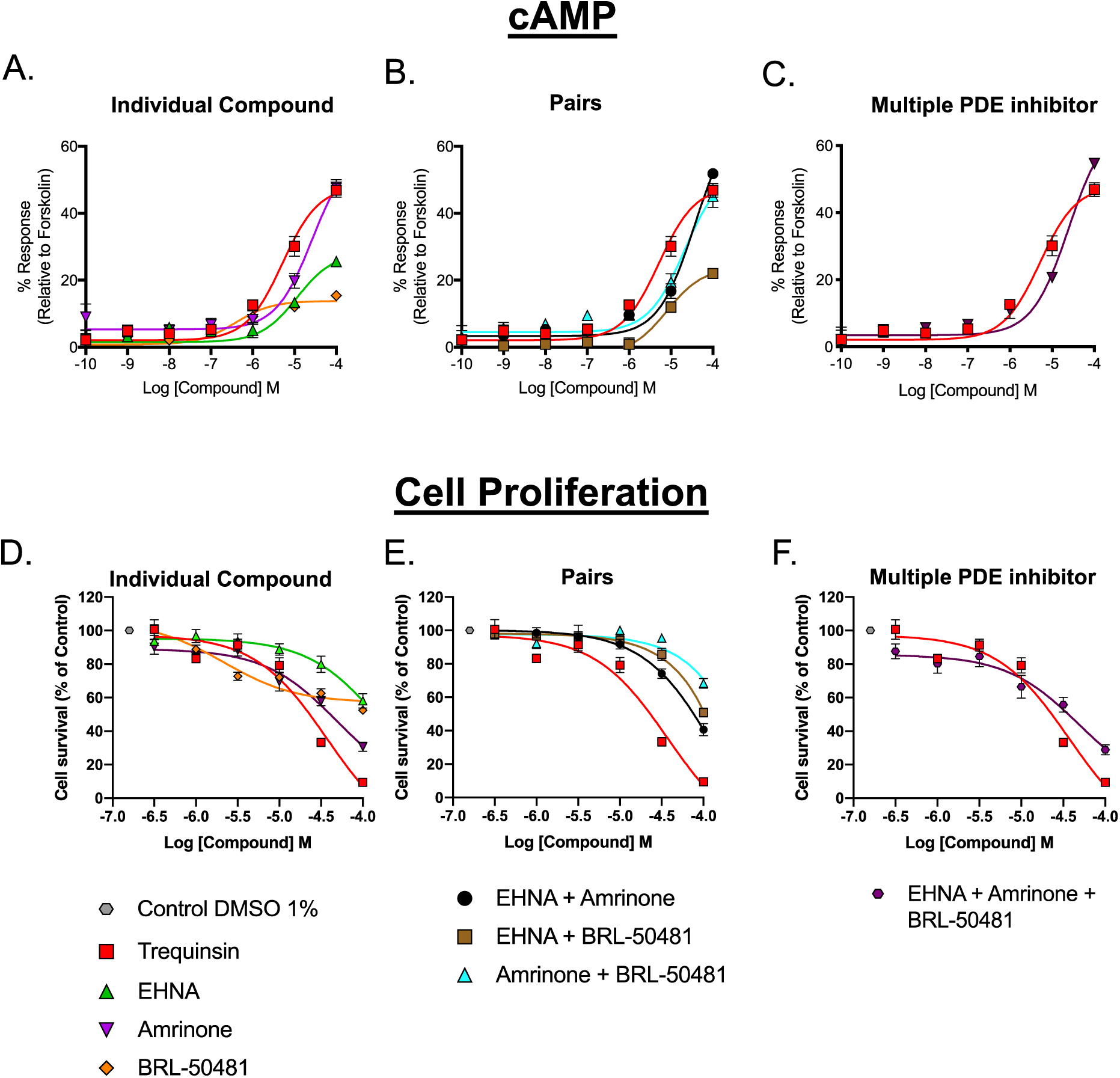
The effect of selective PDE inhibitors, as individual, dual, or multiple treatments, on intracellular cAMP levels and cell proliferation in C6 glioma cells. *A-C.* cAMP accumulation was determined in C6 cells following 30 min stimulation with EHNA, Amrinone or BRL-50481 alone (A), in pairs (B), or combined (C). Data are expressed relative to 100 µM forskolin. *D-F.* Cell survival was determined in C6 cells following 72 hours incubation with EHNA, Amrinone or BRL-50481 alone (D), in pairs (E), or combined (F). Data are expressed as percentage of cell proliferation relative to vehicle from 6-9 data sets. The effect of trequinsin alone is displayed on each graph for comparison. All data are the mean ± SEM of 6–9 individual repeats.

### 3.4 Targeting both AC and PDEs enhances the anti-proliferative effect

Our data suggest that, elevation of cAMP levels either through activation of AC or inhibition of PDEs positively correlated with reduced cell proliferation. Thus, we hypothesised that dual activation of AC and inhibition of PDEs would induce larger suppression in cell growth beyond that of single a target treatment. To test this, we determined the combinatorial effect of forskolin and trequinsin on cAMP accumulation and cell proliferation (Figure 5).

**Figure 5.**
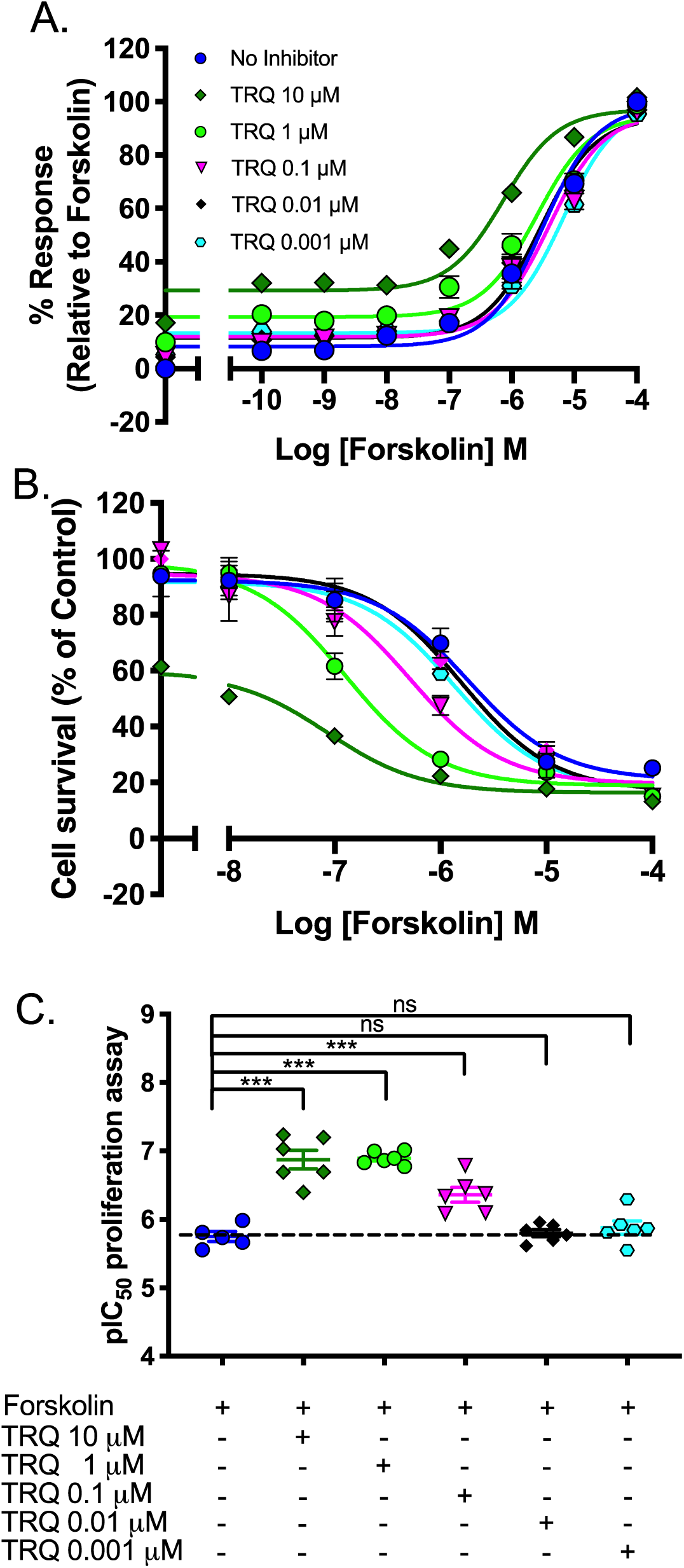
Forskolin and trequinsin act synergistically to increase cAMP accumulation and suppress cell growth. *A*. Concentration-dependent effect of trequinsin upon cAMP accumulation in C6 cells following 30 min stimulation with forskolin. Data are expressed relative to 100 µM forskolin in the absence of trequinsin. *B*. Concentration-dependent inhibitory effect of trequinsin on C6 cell growth following 72-hour incubation with forskolin. Data are expressed as percentage cell proliferation relative to vehicle. *C*. pIC_50_ values for individual cell survival curves for each treatment condition. All data are the mean ± SEM of 6-9 individual repeats. Data were determined as statistically different (ns, not significant; ***, p<0.001) compared to forskolin using one-way ANOVA followed by Dunnett’s *post-hoc* analysis. TRQ – trequinsin.

There was a similar pattern of effects observed upon forskolin and trequinsin co-treatment on both cAMP accumulation and cell proliferation assays. The combination of forskolin and trequinsin significantly enhanced cAMP accumulation and reduced cell growth in a dose-dependent manner compared to forskolin alone in C6 cells (Figure 5A-C). The potency of the forskolin-mediated anti-proliferative effect was enhanced by 13-fold in the presence of 10 μM trequinsin. These data demonstrate synergistic elevation of cAMP by targeting AC and PDEs resulting in greater suppression of C6 cell growth.

### 3.5 The anti-proliferative effect of forskolin is mediated through a PKA-dependent mechanism

Having confirmed the effect of cAMP on cell growth, we next wanted to investigate the involvement of downstream effectors of cAMP such as, protein kinase A (PKA) and exchange protein directly activated by cAMP (Epac) type I and II, as well as the cGMP effector, protein kinase G (PKG), on proliferation of C6 cells (Figure 6A). To do this we utilised a range of small molecule inhibitors KT5720 (PKA), ESI-09 (non-selective Epac), CE3F4 (Epac1), HJC0350 (Epac2), and KT5823 (PKG). Co-treatment of 10 µM KT5720 with forskolin or trequinsin attenuated their anti-proliferative effects on C6 cells (Figure 6B and 6C, by ∼3.16-fold, p<0.005 and trequinsin by ∼1.7-fold, p<0.001). None of the selective or non-selective Epac inhibitors had any effect on trequinsin-stimulated cell growth suppression, although there was an elevation of forskolin-stimulated cell growth suppression with CE3F4 (p<0.01) (Figure 6B). In the presence of KT5720, cAMP is accumulated and provides targets for non-selective PDEs to activate cGMP pathways. Interestingly, cotreatment with KT5823, a PKG inhibitor, enhanced the anti-proliferative effects of forskolin and trequinsin (p<0.001), indicating an involvement of cGMP signalling pathways in cell proliferation. It is possible that accumulation of cGMP in the presence of the PKG inhibitor potentiated cAMP/PKA signalling pathways, through the sequestration of non-selective PDEs, and thereby reduced cell growth. Indeed, further inspection of the effects of GC activators (Figure 1A-C) upon cAMP accumulation and cell proliferation in C6 cells suggests that there is cross-talk between cAMP and cGMP – addition of GC activators induce low levels of cAMP accumulation and limited cell proliferation. These data combined highlight the importance of the homeostasis between cAMP and cGMP and that the anti-proliferative effects of forskolin are mediated through a cAMP/PKA-dependent pathway.

**Figure 6.**
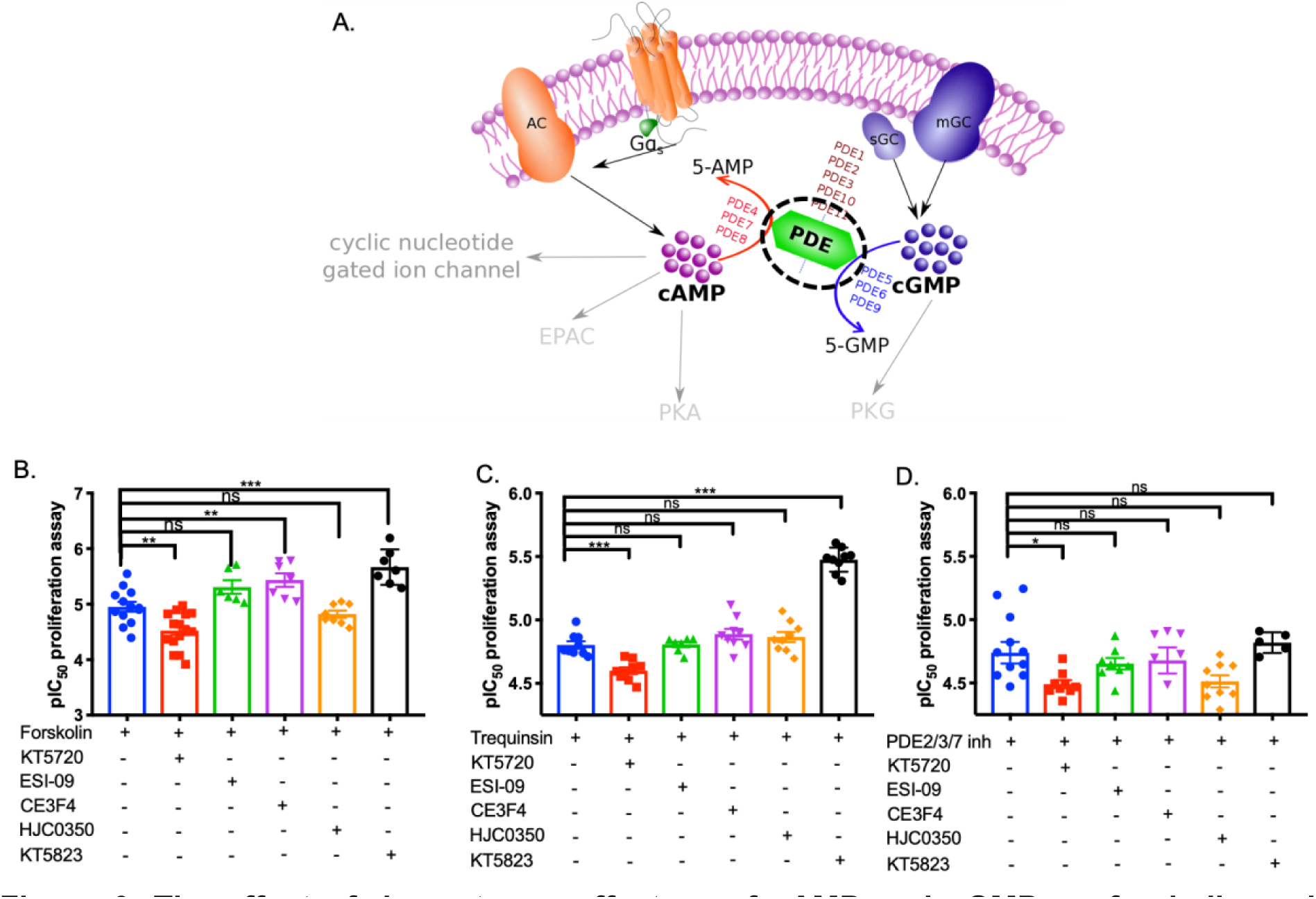
The effect of downstream effectors of cAMP and cGMP on forskolin and trequinsin-mediated cell growth suppression of C6 cells. *A*. Schematic diagram illustrating cAMP and cGMP synthesis, degradation and downstream effectors. *B-D.* Cell survival was determined in C6 cells following 72 hours incubation with forskolin (B), trequinsin (C), or a combination of PDE2,3,7 inhibitors (D) in the presence either KT5720 (10 μM), ESI-09 (10 μM), CE3F4 (10 μM), HJC0350 (10 μM) or KT5823 (10 μM). Data are represented as individual pIC_50_ values for anti-proliferation curves for each treatment condition. Data were determined as statistically different (ns, not significant; *, p< 0.05; **, p<0.01; ***, p<0.001) compared to in the absence of inhibitor using one-way ANOVA followed by Dunnett’s *post-hoc* analysis. KT5720 – PKA inhibitor, ESI-09 - non-selective Epac1/2 inhibitor, CE3F4 – Epac1 inhibitor, HJC0350 - Epac2 inhibitor, KT5823 PKG inhibitor.

### 3.6 Trequinsin may have cAMP-independent action leading to apoptosis but not forskolin and a PDE inhibitor cocktail

In order to delineate the mechanism by which cAMP promotes cell death or inhibit cell growth, we investigated if anti-proliferative effects on C6 cells were related to apoptosis. Early apoptotic events can be detected through the protease activity of caspase-3 and caspase-7 that will eventually degrade proteins pivotal for cell survival. In this study, we quantified positive cell responding to caspase-3 and −7 activity in C6 cells using fluorescence microscopy after treatment with CellEvent ™Caspase-3/-7 green detection kit. Cells were co-stained with Hoechst 33342 and propidium iodide to label all nuclei and dead cells.

Staurosporine is a known pan caspase activator. Quantitative analysis revealed that staurosporine-treated cells were entirely positive for active caspase-3/-7 (Figure 7A). Consistent with these results, activated caspase-3/-7 leads to cell death which was confirmed by propidium iodide staining (Figure 7A). Amongst all other treatments, only 100 µM trequinsin exhibited comparable effects on cell death and caspase activity to that of staurosporine. Treatment with forskolin or the PDE 2,3,7 inhibitor cocktail resulted in less than 10% caspase activity and cell death (Figure 7A). This suggests that 100 µM trequinsin may have toxic, non-cAMP-dependent effects, on C6 cells.

**Figure 7.**
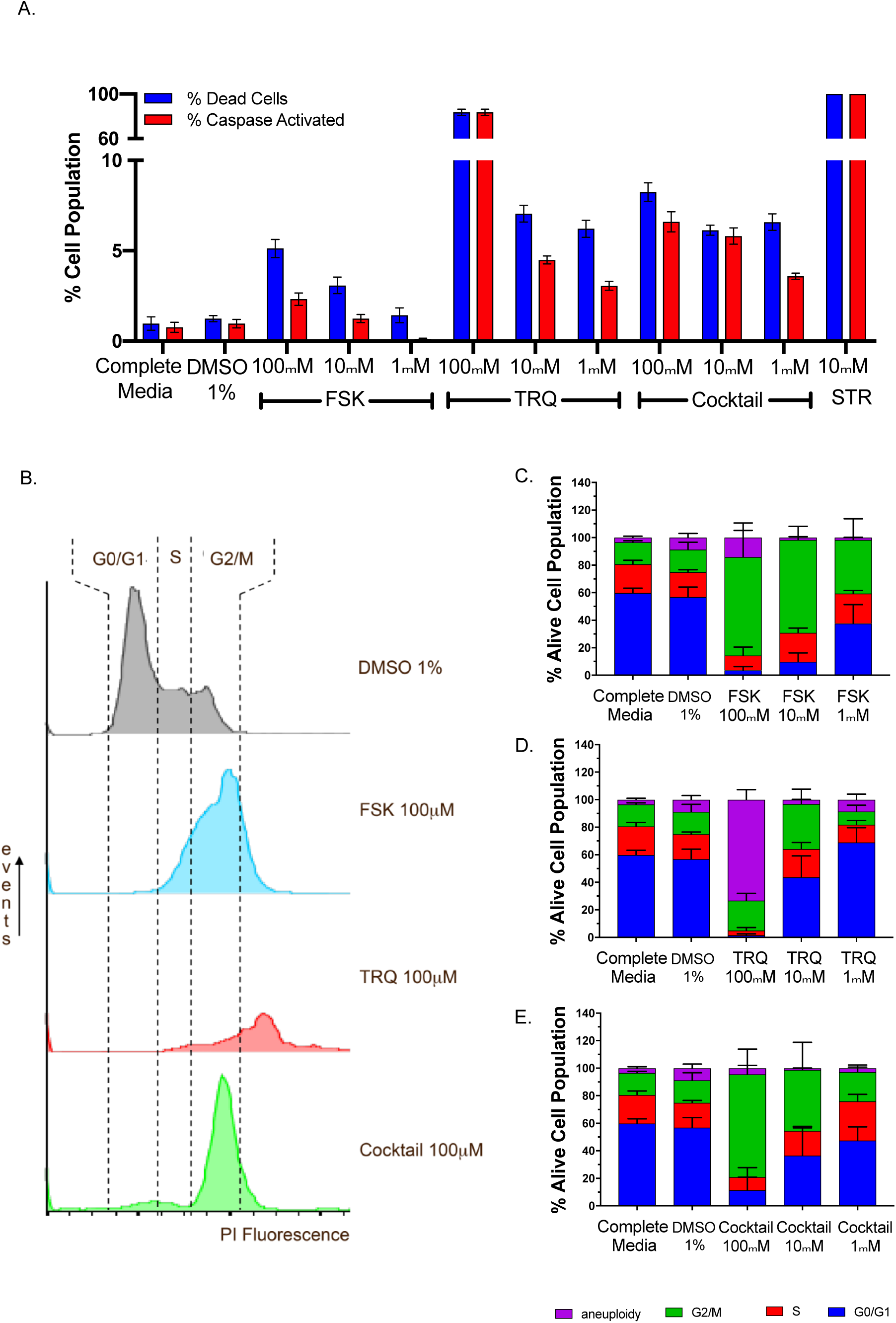
Population of dead cells and activity of caspase-3/7 in C6 cells after 72 h treatment and cell cycle analysis by flow cytometry after PI staining. *A*. Percentage of dead cells determined by staining with propidium iodide. *B.* The percentage of cells with activated caspase-3/7, visualised by CellEvent caspase-3/7. All values are normalised to total cell number in each well. Staurosporine 1 μM was used as a control for apoptotic cell death and cause 100% dead cells. *C-E*. Representative histograms of cells following treatment with forskolin (C), trequinsin (D), or PDE2,3,7 inhibitor cocktail (E) for 72 h. The percentage of cell distribution for each treatment including G_1_, S, G_2_/M, and dead cell population (n= 4-8 individual data). All data are the mean ± SEM of 5 individual repeats. Data were determined as statistically different (*, p<0.05; **, p<0.01; ***, p<0.001) compared to 1% DMSO using a one-way ANOVA followed by Dunnett’s *post hoc* analysis. * p<0.05, ** p<0.005, *** p<0.001. FSK – forskolin, TRQ – trequinsin, STR – staurosporine.

### 3.7 Elevated intracellular cAMP induces growth arrest at the G_2_/M phase of the cell cycle

Having demonstrated that elevation of cAMP inhibits cell proliferation, without inducing extensive apoptotic events, we postulated that this effect arose due to the cell undergoing a cell growth arrest. Thus, we investigated the individual stage of the cell cycle of C6 cells post-treatment with forskolin, trequinsin or the individual PDE2,3,7 inhibitor cocktail by using propidium iodide staining and flow cytometry. Cell cycle analysis showed that forskolin, trequinsin, and the PDE2,3,7 inhibitor cocktail altered cell phase (Figure 7B-E). While there was no significant difference between complete media and DMSO, forskolin or PDE inhibitor cocktail treatment arrested cells predominantly in G_2_/M phase. This indicates that both forskolin and the PDE2,3,7 inhibitor cocktail alters C6 cell cycle by a similar mechanism. In contrast, of the proportion of cells (∼30% that survived treatment with 100 µM trequinsin approximately 70% were aneuploid, with the remaining alive cells arrested in G_2_/M phase. Interestingly, when lower concentrations of trequinsin (<100 µM) were considered, the cell phase profile more closely match that of 10 µM PDE2,3,7 inhibitor cocktail (Figure 7B). This data suggests that 100 µM trequinsin induces a toxic effect on the C6 cells that is most likely independent of its action upon the cAMP-PKA pathway. However, such toxic effects are not observed for the PDE2, PDE3 and PDE7 inhibitor cocktail or forskolin suggesting their actions are dependent upon the cAMP-PKA pathway.

## 4. DISCUSSION

cAMP is a ubiquitous second messenger, which together with cGMP, controls a myriad of physiological responses (Sutherland 1972) including reparative processes. Interestingly, cAMP signalling has been reported to have different effects on cell proliferation; either causing or arresting proliferation, depending on the cell type investigated. These divergent effects of cAMP on the proliferative response are believed to be controlled by several factors including stimulus, the nature of the intracellular cAMP effectors within the cells, the strength of signal, and subcellular compartmentalisation (Insel et al. 2012).

In several types of cancer, malignancy accompanies reduction of cAMP levels. Particularly in brain tumours, suppression of cAMP is associated with glioma genesis compared with non-tumour controls (Warrington et al. 2010, Warrington et al. 2007, Daniel et al. 2016). As cAMP levels are suppressed 4-fold in brain tumours compared to those in normal tissue (Furman and Shulman 1977), we hypothesised that augmenting production of cAMP may restore intracellular signalling and have beneficial antiproliferative effects.

Elevation of intracellular cAMP levels can be achieved through activation of Gα_s_-protein coupled receptors, stimulation of AC or inhibition of PDEs. While forskolin directly stimulates AC to increase production of cAMP, PDE inhibitors prevent cAMP breakdown and, PTX and CTX modify Gα_i_ and Gα_s_, respectively, to increase Gα_s_ activation of AC. However, neither PTX nor CTX alone are sufficient to induce cAMP-mediated growth suppression, possibly due to transient activation of cAMP downstream cascade or receptor desensitisation. Conversely, treatment with forskolin or selected PDE inhibitors induce a greater elevation of intracellular cAMP levels and potently suppresses cell growth. Taken together, our results suggest that only direct activation of AC or inhibition of PDEs elevates global cAMP levels sufficiently to attenuate cell growth and/or trigger apoptosis.

With the exception of PDE4C, PDE8B, and PDE11A, almost all PDE isoenzymes are present in C6 glioma cells. C6 cells expressed a greater number of PDEs and to a greater extent compared to ST14A, a healthy neuronal-derived cell line. Differential expression of PDEs between glioma and normal brain cells supports the fact that glioma has a compensatory mechanism to maintain low levels of cAMP. In this study, we have provided evidence that augmenting cAMP levels in C6 cells positively correlates with anti-proliferative effects.

Inhibitors of cAMP-specific or cGMP-specific PDEs showed minimal effects on cell growth, except for ibudilast (a cAMP-specific PDE4 inhibitor). Inhibition of PDE1, PDE2, PDE3, PDE4, and PDE7 resulted in better modulation of cAMP levels and cell growth. These PDEs, with the exception of PDE4 and PDE7, hydrolyse both cAMP and cGMP. It appears that an imbalance between intracellular cAMP and cGMP levels mediated by PDEs may have consequences on downstream intracellular signalling cascades that is ultimately responsible for modulating cell growth. Our hypothesis is supported by the results of a series of so called ‘rescue experiments’, whereby inhibition of PKA, but not PKG, in the presence of forskolin or trequinsin attenuated anti-proliferative effects. PKG inhibition increased anti-proliferative effects and thus may disturb the homeostasis of cAMP and cGMP by providing more targets for activating cAMP pathways. Interestingly, only PDE inhibitors that work on dual substrate PDEs (which include PDE2 and PDE3) exhibiting better efficacy both for cAMP accumulation and cell proliferation assay. All these results provide an insight that disturbance of cAMP-cGMP levels trigger modulatory effects leading to augmentation of cAMP/PKA signalling pathways.

The most potent anti-proliferative effects were observed in trequinsin-treated cells. Trequinsin is commonly known as an ultrapotent PDE3 inhibitor, although it has also been shown to have activity against PDE2 and PDE7 (Rickles et al. 2010). There have, however, been no studies into inhibition of PDE2 and PDE7 by trequinsin. We sought to elucidate the mechanism by which trequinsin mediates its antiproliferative effects. A combination of inhibitors of PDE2, PDE3, and PDE7 had a similar magnitude of effect to that of trequinsin, whilst inhibition of single or dual PDEs failed to mimic the effect. This implies that trequinsin may indeed inhibit all three PDEs. In order to enhance antiproliferative activity further, the selectivity of individual compounds towards each PDEs needs to be improved.

We identified that multitarget enhancement of cAMP signalling elevated anti-proliferative effects of a glioma cell model. Increasing cAMP levels through activation of AC and multiple PDE inhibition demonstrated synergistic cell growth suppression. Our results highlight that the dual effect of AC activation and PDE inhibition triggers a substantial increase in intracellular cAMP levels compared to single target treatments resulting in more potent anti-proliferative effect.

By applying several tools that target downstream effectors of cAMP and cGMP, we determined that forskolin and trequinsin mediate inhibition of cell growth through the activation of PKA. Suppression of C6 cell proliferation by forskolin and trequinsin was significantly attenuated by PKA inhibition, whilst selective (CE3F4 against Epac1 and HJC0350 against Epac2) and non-selective (ESI-09) inhibition of Epac or PKG failed to reduce inhibition of proliferation. Interestingly, inhibition of Epac1 by CE3F4 significantly potentiated the antiproliferative effect of forskolin on C6 cells. This is most likely due to cAMP produced by forskolin effectively being diverted away from Epac1 and towards the PKA pathway.

Although trequinsin has a comparable potency to that of forskolin for elevating intracellular cAMP levels and supressing cell proliferation, at high concentrations (100 μM) trequinsin induced substantial cell death of C6 cells and ST14A cells. This implies that at such high concentrations trequinsin binds to other non-PDE proteins that results in direct activation of caspase-3/7 to trigger cell death. Indeed, assessment of caspase-3/7 activity revealed that 100 μM trequinsin triggered cell death predominantly by activating caspase-3/7. However, we have demonstrated synergism between activation of AC (forskolin) and inhibition of PDE2/3/7 (trequinsin) that significantly abrogates cell growth. Thus, the dose of each compound can be reduced to potentially prevent any toxic side effects of trequinsin.

Alternatively, combining individual selective inhibitors against PDE2,3,7 together showed similar efficacy to that of trequinsin but without significant toxicity in glioma cells. The inhibitor cocktail shows a higher number of cells with active caspase-3/7 activity compared to forskolin, possibly due to the availability of PDEs, which have been shown to be caspase substrates (Lerner and Epstein 2006). Both forskolin and the inhibitor cocktail trapped cells in G_2_/M phase, which may lead to the loss of essential cellular components that are required for replication. Conversely, the small population of cells that survived treatment with 100 μM trequinsin were aneuploid.

In this study, we have used a chemical biology approach to demonstrate that cAMP inhibits growth of a glioma cells. Anti-proliferative effects of forskolin are mediated by elevating cAMP levels leading to activation of PKA and arrest of cell sin the G_2_/M phase of the cell cycle. In comparison, multiple inhibition of PDEs by trequinsin not only inhibits cell growth via the cAMP/PKA cascade, but also triggers cell death through caspase-3/-7 activation. Concomitant targeting of both AC and PDEs to synergistically elevate intracellular cAMP levels within glioma cells. Due to possible side effects of trequinsin, a cocktail of individual PDE2, PDE3, and PDE7 inhibitors can be used as an alternative to trequinsin to obtain similar functional effects without any toxicity. This study offers insight to identify new therapeutic approaches which have potential beneficial effects against glioma/glioblastoma.

## SIGNIFICANCE

The suppression of intracellular concentrations of cAMP have been reported to be associated with malignancy. Thereby, restoration of cAMP levels is a potential method to suppress cell growth. Previously, several approaches have been tested using cAMP analogues. These, however displayed toxicity and poor selectivity. Here, using C6 cells as a model of glioma, we have found that phosphodiesterases (PDEs) – enzymes that degrade both cAMP and cGMP - are more highly expressed in C6 cells than in a model of healthy striatal-derived cells. Application of small molecule pan-activators of AC or selective PDE inhibitors, enabled us to assess the ability of each compound to modulate cAMP and cell proliferation. Here, we demonstrate that chemical enhancement of cAMP levels directly correlates with anti-proliferative actions on C6 cells. Interestingly, trequinsin, a small molecule compound which has been reported to antagonise multiple PDEs, displayed the most potent activity among all PDE inhibitors tested. We also highlight that combining the small molecules, forskolin and trequinsin, synergistically enhances cAMP accumulation, beyond that of the individual compounds, leading to improved potency for inhibiting cell proliferation. Further, we demonstrate, by the use of chemical inhibitors of downstream effectors of cAMP, that the enhanced cAMP levels are directly responsible for decreased cell proliferation (cells become arrested in the G_2_/M phase of the cell cycle). At high concentrations of trequinsin (>10 µM) we observed cAMP-independent cell toxicity, thus limiting its use as a monotherapy. However, through generating a PDE2,3,7 inhibitor cocktail we could achieve cell cycle arrest without any toxic effects. We suggest that the data presented here validates the use of small molecule compounds that enhance the intracellular concentrations of cAMP as inhibitors of cell proliferation.

## AUTHOR CONTRIBUTIONS

DS, LK, FS, DB, GL conceived and designed the research; DS, HP, IW performed the experiments; DS, MTH, TR, and GL analyzed data; DS, MH, HYY and GL wrote manuscript, DB, LK, TR, MTH revised and edited the manuscript.

## ACKNOWLEDGEMENTS

Authors acknowledge the support of Endowment Fund for education from Ministry of Finance Republic of Indonesia (DS), a BBSRC Flexible Talent Mobility Account scheme award (GL and LK), BBSRC Doctoral Training Partnership BB/JO1454/1 (MH), Rosetrees foundation (to HYY and GL), and the Brain Tumour Charity (UK) grant GN-000429 (LK, FS, DB). HYY is also supported by an international scholarship from the Cambridge Trust. We thank Sampurna Chakrabarty for support throughout preparation of this article. Finally, we would like to thank Colin Taylor for the C6 cell line.

## DECLARATIONS

DB and LK are current employees of *IOTA Pharmaceuticals Ltd*; FS is a former employee.

