## Supplemental figures for "Elevated intracellular cAMP concentration mediates growth suppression in glioma cells"

**Supplemental Information**

**Figure 1. Individual dose-response curve of each PDE inhibitor (PDE1 – PDE5 inhibitor) upon cAMP accumulation on both C6 and ST14A cells.**

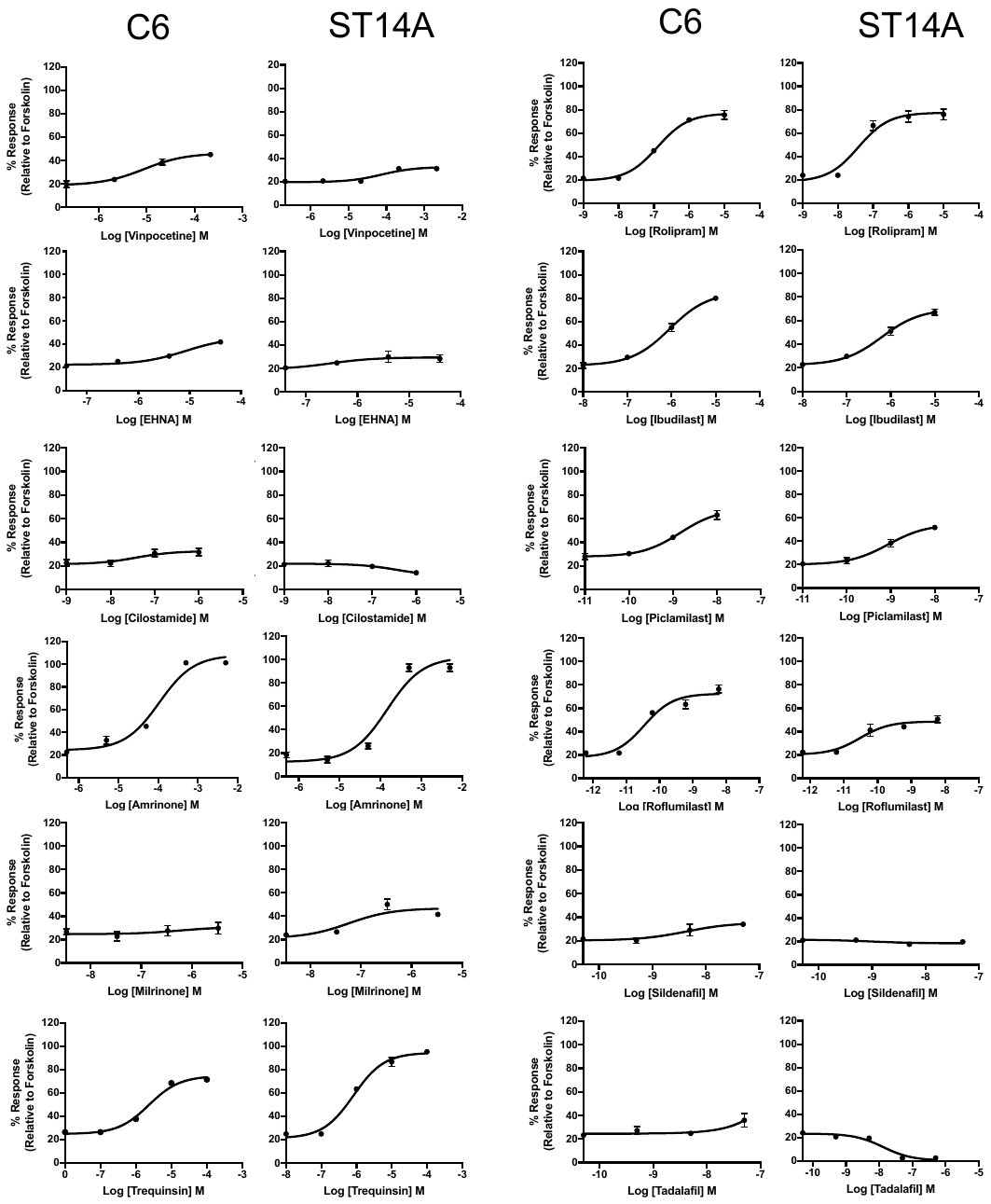

**Figure 2. Individual dose-response curve of each PDE inhibitor (PDE5 – PDE11 inhibitor) and forskolin upon cAMP accumulation on both C6 and ST14A cells.**

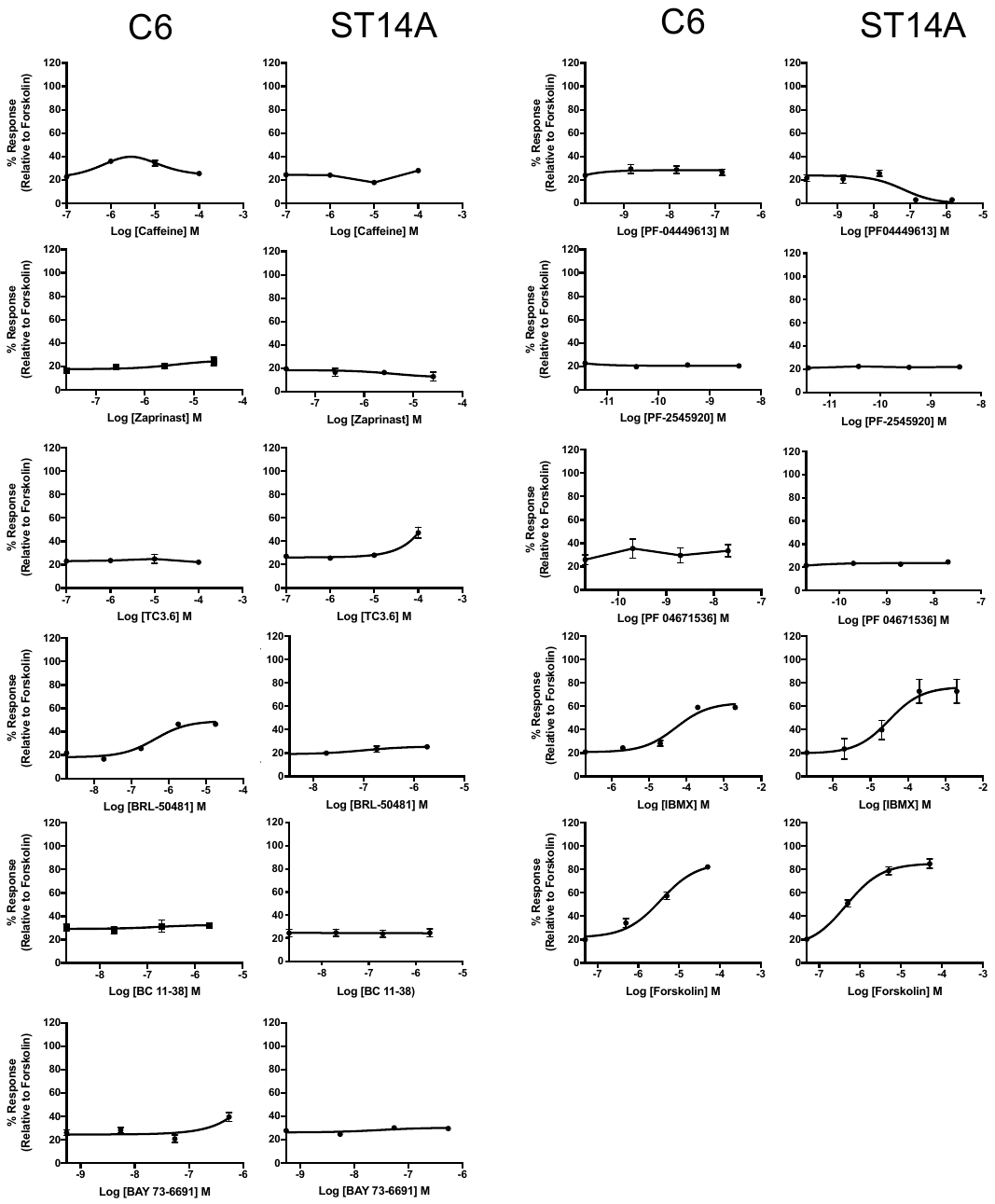

**Figure 3. Individual dose-response curve of each PDE inhibitor (PDE1 – PDE5 inhibitor) upon cell proliferation on both C6 and ST14A cells.**

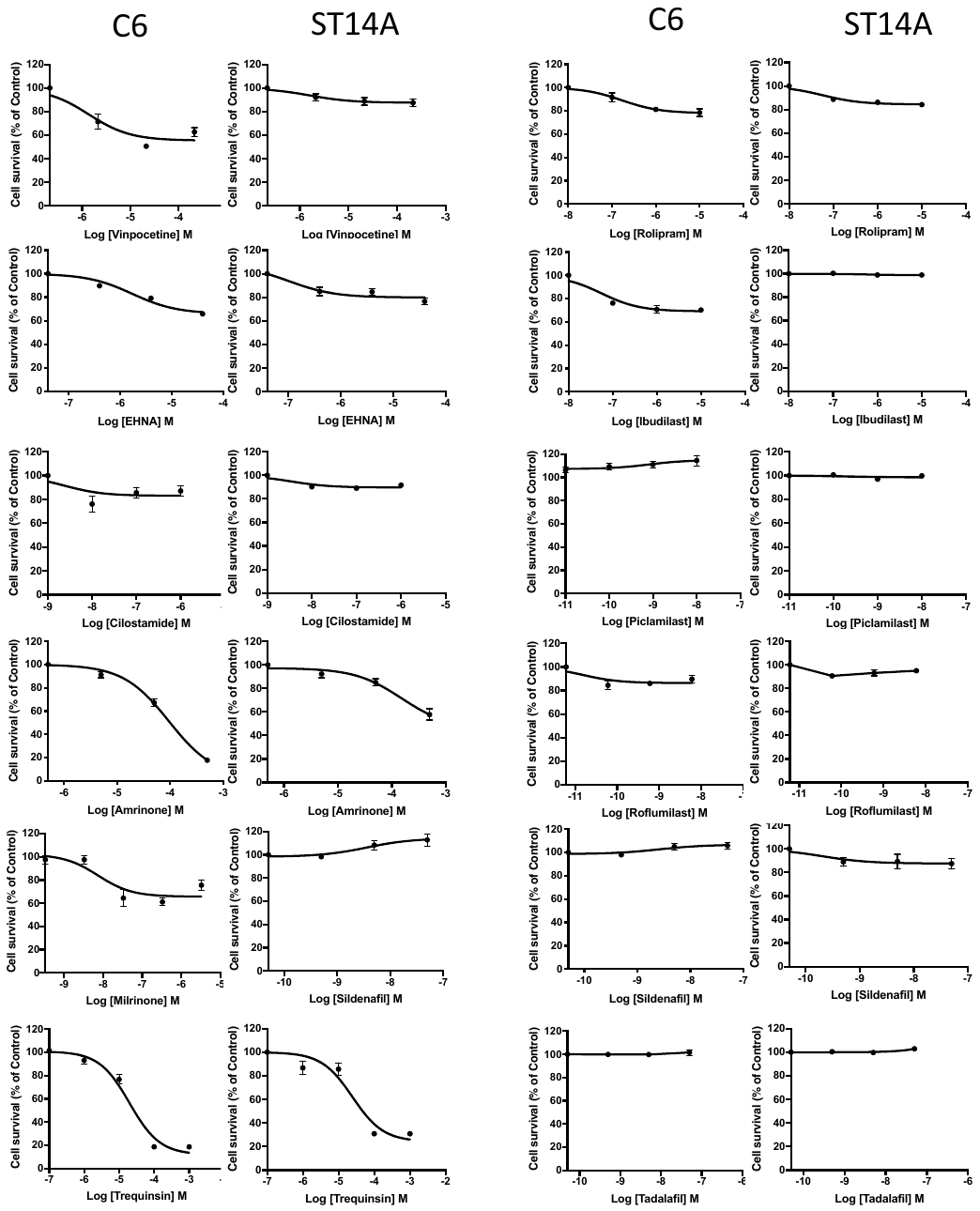

**Figure 4. Individual dose-response curve of each PDE inhibitor (PDE5 – PDE11 inhibitor) and forskolin upon cell proliferation on both C6 and ST14A cells.**

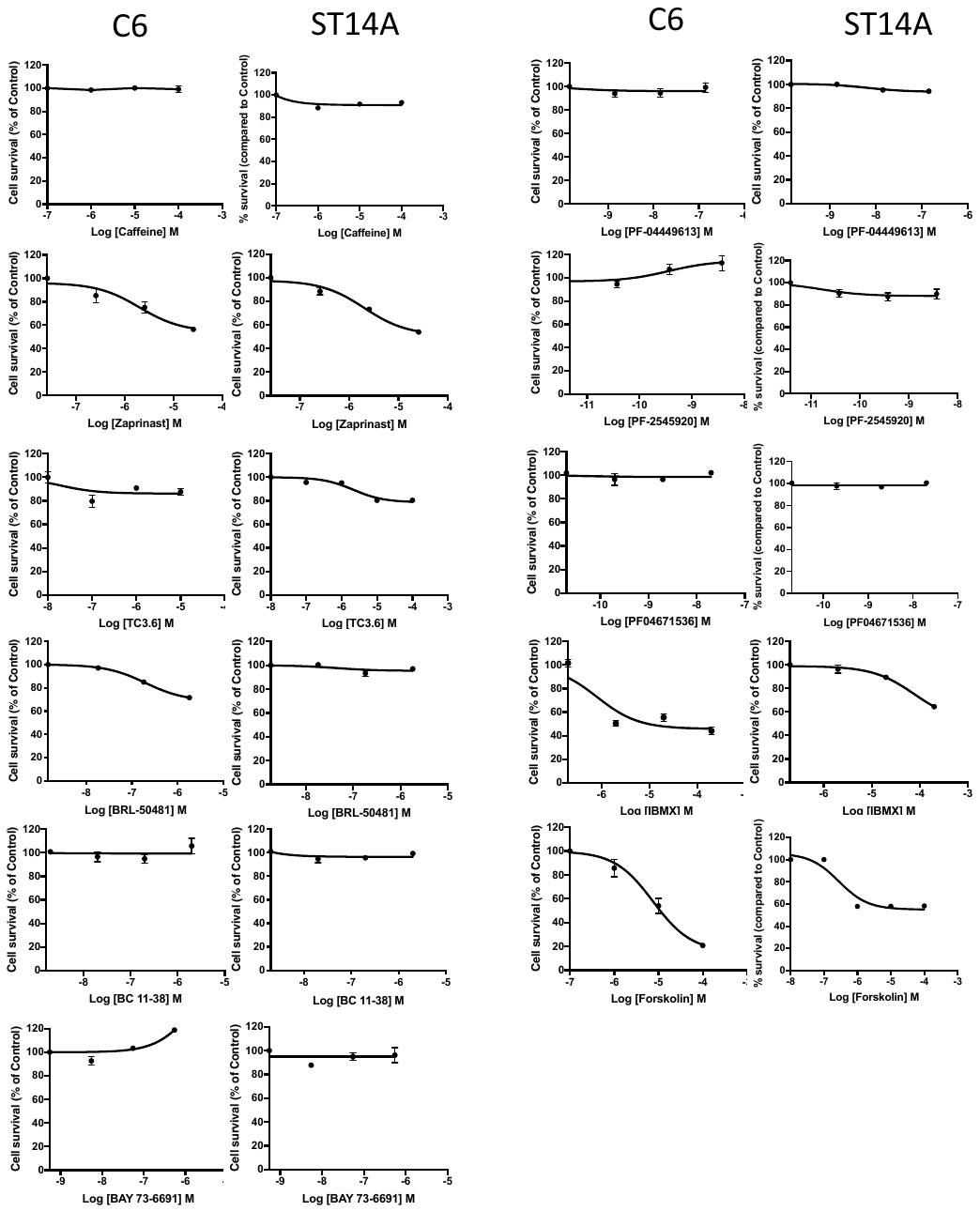
